# bHLH heterodimer complex variations shape meristems in *Arabidopsis thaliana* by affecting target gene specificity

**DOI:** 10.1101/2022.06.02.494582

**Authors:** Eliana Mor, Markéta Pernisová, Max Minne, Guillaume Cerutti, Dagmar Ripper, Jonah Nolf, Jennifer Andres, Laura Ragni, Matias D. Zurbriggen, Bert De Rybel, Teva Vernoux

**Affiliations:** Ghent University, Department of Plant Biotechnology and Bioinformatics, Technologiepark 71, 9052 Ghent, Belgium; VIB Centre for Plant Systems Biology, Technologiepark 71, 9052 Ghent, Belgium; Laboratoire Reproduction et Développement des Plantes, Univ Lyon, ENS de Lyon, CNRS, INRAE, INRIA, 69342 Lyon, France; Functional Genomics and Proteomics, National Centre for Biomolecular Research, Faculty of Science, and Plant Sciences Core Facility, Mendel Centre for Plant Genomics and Proteomics, Central European Institute of Technology (CEITEC), Masaryk University, 62500 Brno, Czechia; ZMBP-Center for Plant Molecular Biology, University of Tübingen, Auf der Morgenstelle 32, D-72076 Tübingen, Germany; Institute of Synthetic Biology and CEPLAS, Heinrich-Heine-Universität Düsseldorf, Universitätsstrasse 1, 40225 Düsseldorf, Germany

## Abstract

The main regions of cell proliferation in plants are the root and shoot apical meristems during primary growth and the vascular cambia as lateral meristems during secondary thickening. A number of unique regulators have been described in each of these meristems, suggesting that these different meristems might have independently evolved dedicated transcriptional networks to balance cell proliferation. Here, we show that the basic Helix Loop Helix (bHLH) transcription factor complexes formed by TARGET OF MONOPTEROS5 (TMO5), LONESOME HIGHWAY (LHW) and their close homologs are broadly expressed throughout plant development and operate as general regulators of cell proliferation in all meristems. Yet, genetic and expression analyses indicate that these complexes have specific functions in distinct meristems mediated by heterodimer complex variations between members of the TMO5 and LHW subclades. We determine that this is primarily due to their expression domains limiting the possible combinations of heterodimer complexes within a certain meristem, and to a certain extent to the absence of some members in a given meristem. We further demonstrate target gene specificity for heterodimer complexes, suggesting that spatial differences in transcriptional responses through heterodimer diversification allow a common bHLH heterodimer complex module to contribute to the control of cell proliferation in multiple meristems.

## INTRODUCTION

Post-embryonic plant growth and development is driven by the activity of three main pools of pluripotent stem cells contained in zones called meristems. These are tightly regulated to divide and differentiate into specific cell types and form new organs. The root apical meristem (RAM) is located at the growing root tip, laid down during embryogenesis and responsible for a formation and primary growth of below-ground organs. At the other extremity, an activity of stem cells in the shoot apical meristem (SAM) is responsible for aerial organ development (Wang et al., 2018). While the apical meristems (RAM and SAM) give rise to the primary plant body, plants use the third pool of proliferating cells located in lateral meristems (LM) to support secondary growth leading to an increase in root and stem girth or thickness (Ragni and Greb, 2018). These meristems represent vascular and cork cambia (Etchells and Turner, 2010; Serra et al., 2022). Meristem activity is essential for growth and development and thus needs to be tightly controlled to ensure optimal growth depending on the environmental conditions and to avoid excessive cell proliferation (Motte et al., 2019).

Several key regulators, including transcription factors (TFs), and ligand-receptor complexes have been identified, which contribute to this intricate regulation of each of these meristem regions (Shimotohno and Scheres, 2019). For example, the CLAVATA3 (CLV3)-CLV1-WUSCHEL (WUS) negative feedback loop is the central genetic mechanism that coordinates stem cell proliferation with differentiation in the SAM. Perturbation of this regulatory network leads to phenotypes ranging from a loss of the meristem to a massive over-proliferation of meristematic cells (Clark et al., 1993, 1995; Laux et al., 1996; Gaillochet et al., 2015). Regulation of the LM that contributes the most to radial growth, called the vascular cambium, occurs via CLAVATA3-LIKE/ESR-RELATED 41 (CLE41) peptides produced in the phloem and perceived in the cambium by the PHLOEM INTERCALATED WITH XYLEM (PXY)/ TDIF RECEPTOR (TDR) receptor. Through activation of the direct targets of the complex, *WUSCHEL RELATED HOMEOBOX 4* (*WOX4*) and *WOX14*, this pathway regulates cell division and vascular patterning (Fisher and Turner, 2007; Suer et al., 2011; Etchells et al., 2013). In the RAM, the peptide-receptor kinases complex formed by CLE40-CLV1-ACT DOMAIN REPEAT 4 (ACR4) controls *WOX5* expression and activity, thereby orchestrating stem cell maintenance and balancing the differentiation activity (Stahl et al., 2013; Berckmans et al., 2020). The DNA binding with One Finger (DOF)-type TFs have also been shown to control cell division orientation and proliferation in the vascular cells in the RAM in a cytokinin dependent manner (Miyashima et al., 2019; Smet et al., 2019). These act downstream of the basic helix-loop-helix (bHLH) TFs complex formed by TARGET OF MONOPTEROS5 (TMO5) and LONESOME HIGHWAY (LHW) (De Rybel et al., 2013; Ohashi-Ito et al., 2013a, 2013b, 2014).

So far, studies have thus focused on factors that seem to be almost exclusively specific to one of the three meristem regions. While dedicated regulatory networks are likely required in different meristems, the alternative possibility remains that we are simply yet to uncover common factors required to regulate proliferation in all meristems. The TMO5/LHW bHLH heterodimer complex is a good candidate to function in multiple meristems as individual members have been shown to be broadly expressed in vascular tissues throughout plant development (De Rybel et al., 2013; Ohashi-Ito et al., 2013a, 2013b, 2014). Moreover, bHLH dimers are well suited to allow diversification in functions by using three main parameters: spatiotemporal expression patterns, DNA binding specificity, and dimerization properties (Grove et al., 2009). Indeed, bHLH TFs display a variety of expression patterns, where the overlap can define their sites of actions in space and time (Qian et al., 2021; Hao et al., 2021). DNA binding specificity is dictated by a highly conserved signature of amino acid motif that forms the basic DNA binding regions and shows significant variations in the bHLH family (Massari and Murre, 2000). Finally, specificity in dimerization properties was highlighted as a determining factor for the majority of bHLH proteins (Grove et al., 2009).

TMO5 acts downstream the auxin-dependent transcription factor MONOPTEROS/AUXIN RESPONSE FACTOR5 (MP/ARF5) (Schlereth et al., 2010). TMO5 has three homologs, TMO5 LIKE1-3 (T5L1-3), all showing a similar expression pattern restricted to the xylem in the RAM (De Rybel et al., 2013). Loss-of-function of TMO5 and its closest homolog T5L1 leads to a reduced vascular cell number compared to wild-type (WT) and a monarch patterning defect with only one pole of phloem and xylem, compared to the diarch WT phenotype (De Rybel et al., 2013). Higher order mutants enhance the severity of these phenotypes, suggesting they are redundant family members (De Rybel et al., 2013). Similarly, LHW also has three homologs, LHW LIKE1-3 (LL1-3). Although LHW and its homologs have a broader expression pattern in the RAM (Ohashi-Ito and Bergmann, 2007; De Rybel et al., 2013; Ohashi-Ito et al., 2013a, 2013b), defects in LHW lead to identical phenotypes as the *tmo5 t5l1* double mutant (De Rybel et al., 2013; Ohashi-Ito et al., 2013a, 2013b). Moreover, higher order mutants increase the severity of the phenotypes, indicating that their function is dose-dependent (Ohashi-Ito and Bergmann, 2007; De Rybel et al., 2013). Combined misexpression of both TMO5 and LHW factors triggers ectopic periclinal and radial cell divisions throughout the RAM, suggesting that they function as part of an obligate heterodimer complex (Ohashi-Ito and Bergmann, 2007; De Rybel et al., 2013; Ohashi-Ito et al., 2013a, 2013b; Smet et al., 2019). Indeed, members of the TMO5 and LHW subclades, which overlap in expression in the young xylem cells of the primary RAM, interact and form heterodimers (De Rybel et al., 2013; Ohashi-Ito et al., 2013a, 2013b, 2014). The TMO5/LHW complex directly activates expression of *LONELY GUY3* (*LOG3*), *LOG4* and *BETA-GLUCOSIDASE44* in the xylem cells, leading to higher levels of active cytokinins by increasing biosynthesis (LOG3/4) and deconjugation (BGLU44) (De Rybel et al., 2014; Ohashi-Ito et al., 2014; Yang et al., 2021). Cytokinins are thought to diffuse to the neighbouring procambium and phloem cells where they trigger divisions. This activity is balanced by CYTOKININ OXIDASE3, which is induced by *SHORT ROOT*, itself a direct TMO5/LHW target gene (Yang et al., 2021).

Here, we show that the TMO5/LHW complex activity is not restricted only to the primary root meristem region but is more broadly required for normal development of RAM, SAM and the vascular cambium as LM. The required diversity to control these differently organized meristems is obtained by differences in expression domains and heterodimer complex variations between members of the TMO5 and LHW subclades, which lead to target gene specificity.

## RESULTS

### TMO5/LHW function is not restricted to primary root development

To establish if the function of TMO5 and LHW clade members is restricted to the RAM or whether they play a broader role during plant development, we first explored the effect of altered heterodimer levels in the SAM and in the vascular cambium during root secondary growth, using existing higher order mutants (*tmo5 t5l1* double, *tmo5 t5l1 t5l3* triple, and *lhw ll1* double mutants) (De Rybel et al., 2013; Ohashi-Ito et al., 2013a) and a constitutive misexpression line (*ProRPS5A::TMO5* x *ProRPS5A::LHW*) (De Rybel et al., 2013) in comparison to wild type Col-0. Given that vascular cell numbers are not easily quantified in the SAM and that the capacity of TMO5 and LHW to induce cell division is not limited to vascular cells (De Rybel et al., 2013), we measured the SAM area as a read-out for a possible effect on cell proliferation activity. We found significant changes in the SAM area in all the lines compared to control (**Figure 1 A-E and K**) (see example of SAM surface analysis in **Supplemental Figure S1A**). The misexpression line, the *tmo5 t5l1 t5l3* and *lhw ll1* mutants had a smaller SAM while *tmo5 t5l1* had a slightly but significantly bigger SAM area. Changes in SAM size were only partial due to changes in cell size (**Figure 1L**) and cell number (**Supplemental Figure S1B**), indeed suggesting a role of these genes in the regulation of cell proliferation throughout the SAM which is more complex compared to the RAM. Similar to the effects observed in the primary RAM, the number of vascular cell files was reduced during root secondary growth in a dose-dependent manner in the loss-of-function mutant lines and increased in the misexpression line (**Figure 1 F-J and M)**. In summary, these results suggest that the activity of TMO5/LHW and some of their homologs might not be restricted only to control cell proliferation in the primary RAM but in other meristems as well.

**Figure 1.**
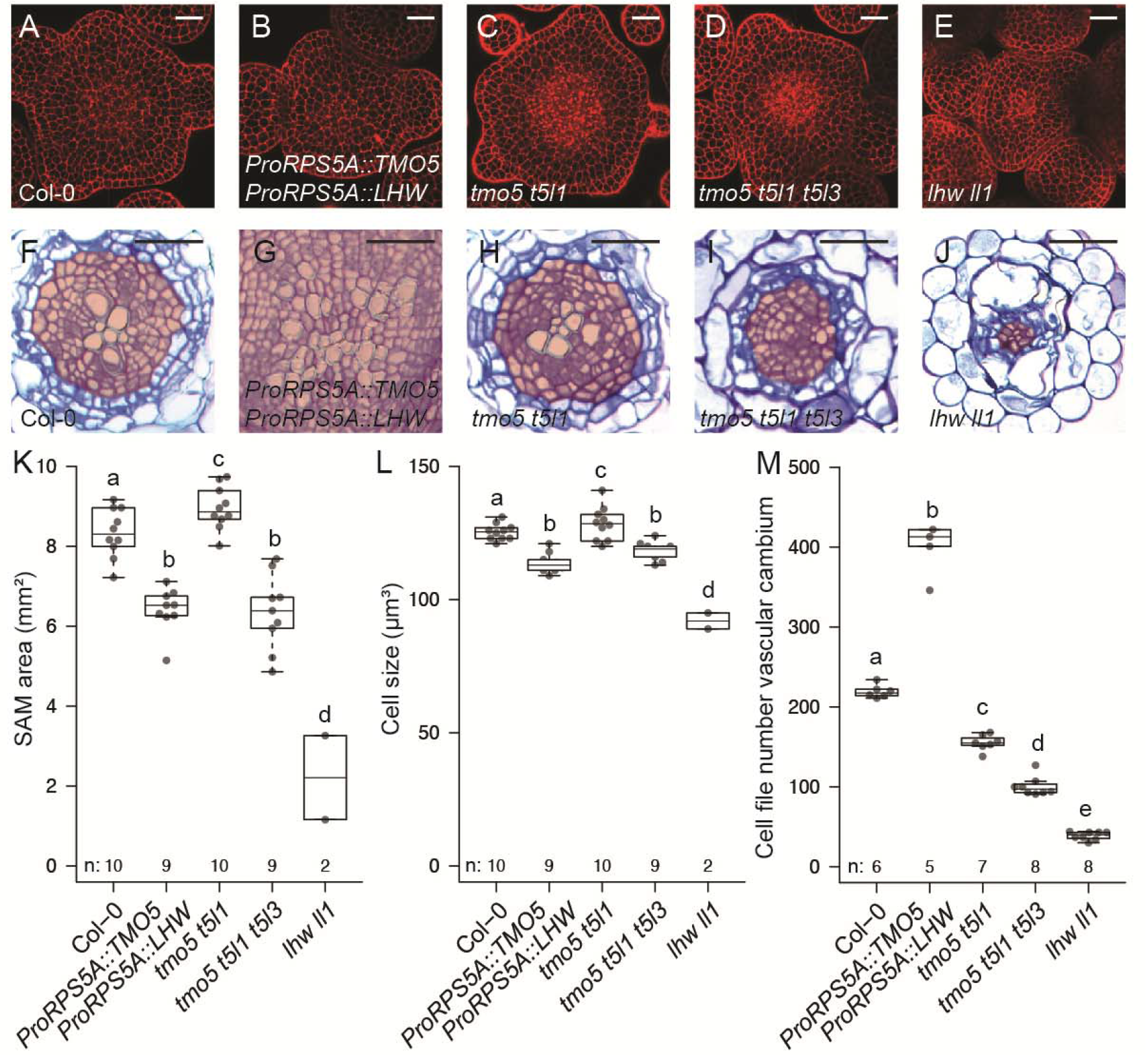
TMO5/LHW function is not restricted to primary root development. Cross sections of shoot apical meristems and roots undergoing secondary growth (uppermost part of the root) of 10-day-old seedlings of (A, F) wild type Col-0; (B, G) *ProRPS5A*::TMO5 x *ProRPS5A*::LHW; (C, H) *tmo5 t5l1*; (D, I) *tmo5 t5l1 t5l3* and (E, J) *lhw ll1*. (K) Determination of shoot apical meristem area and (L) cell size in L1 layer, and (M) quantification of vascular cell files (within but excluding the pericycle, red coloured area in F-J) number of root cross sections. Lowercase letters in charts indicate significantly different groups as determined using a one-way ANOVA with post-hoc Tukey HSD testing (p ≤ 0.05). Scale bars: (A-E) 20 µm; (F-J) 100 µm.

### TMO5/LHW is required and sufficient for root secondary growth

Our results suggest that the TMO5/LHW pathway is conserved in function in primary meristems and during root secondary growth. However, it remains possible that the observed effects during secondary growth are a consequence of the persistent lack or overexpression of these factors during primary growth. Thus, to investigate if TMO5 is sufficient to trigger vascular proliferation during secondary growth specifically, TMO5 was exclusively expressed during this developmental stage by introducing a dexamethasone (DEX) inducible *ProRPS5A::TMO5-GR* rescue construct into the *tmo5 t5l1 t5l3* triple mutant. When grown on medium supplemented with 10 μM DEX, the *ProRPS5A::TMO5-GR* construct introduced in the triple mutant can rescue the proliferation defect in the *tmo5 t5l1 t5l3* triple mutant to an almost non-phenotypical (*t5l1 t5l3* double mutant) situation (**Figure 2 A, B and I**) (De Rybel et al., 2013). This inducible rescue system was next used as a tool to investigate if TMO5 expression during root secondary growth is sufficient to trigger vascular cell proliferation. *tmo5 t5l1 t5l3* mutants with and without the inducible *ProRPS5A::TMO5-GR* rescue construct were grown on mock medium for 5 days and then transferred onto inducing medium supplemented with 10 μM DEX for another 5 days. Again, the number of vascular cell files in the root undergoing secondary growth was quantified. *tmo5 t5l1 t5l3* mutants carrying the *ProRPS5A::TMO5-GR* rescue construct showed a significant increase in vascular cells numbers compared with the triple mutant without the rescue construct (**Figure 2 C, D and I**). This indicates that induction of TMO5 specifically during secondary growth is sufficient to trigger periclinal division leading to radial expansion. Next, we made use of the same rescue system to investigate if the TMO5/LHW pathway is required for secondary growth. Seedlings were first grown on inducible DEX medium for 5 days and then transferred onto mock medium for an additional 10 days. This timing was chosen as the effect of the initial 5 days DEX treatment persists for several days after transfer to mock medium **(Figure 2 E, F and I)**. After the transfer to mock medium and an additional 10 days of growth, no significant difference could be observed between the number of vascular cell files of the triple mutant compared to the triple mutant carrying the *ProRPS5A::TMO5-GR* rescue construct (**Figure 2 G-I**), indicating that TMO5 presence during primary growth is sufficient to initiate secondary growth but not to maintain it. These results therefore suggest that the TMO5/LHW pathway is both sufficient and required to allow vascular proliferation during root secondary growth.

**Figure 2.**
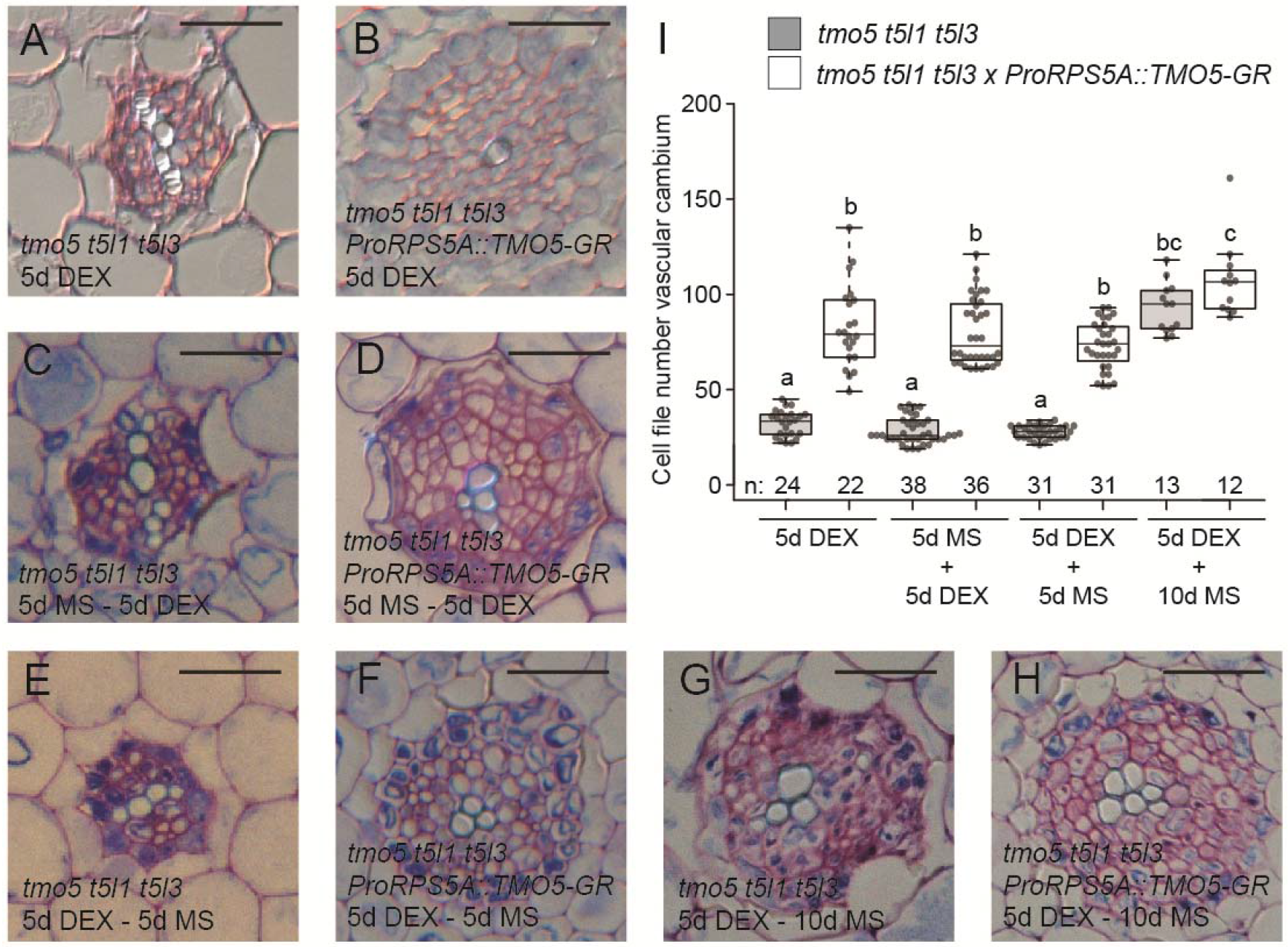
TMO5/LHW is required and sufficient for root secondary growth. Cross sections of roots undergoing secondary growth (upper most part of the root) of *tmo5 t5l1 t5l3* and *tmo5 t5l1 t5l3* with *ProRPS5A*::TMO5-GR seedlings grown either (A, B) 5 days on medium supplemented with 10 µM dexamethasone (DEX); (C, D) 5 days on mock medium (MS) and then transferred for additional 5 days to DEX; (E, F) 5 days on DEX and then transferred for 5 or (G, H) 10 days to MS medium. (I) Quantification shows vascular cell files number of cross sections. Lowercase letters in chart indicate significantly different groups as determined using a one-way ANOVA with post-hoc Tukey HSD testing (p ≤ 0.05). Scale bars: 100 µm.

### TMO5 and LHW clade members show overlapping expression in distinct meristems

Although the redundant role of both TMO5 and LHW subclade members in vascular proliferation of the primary RAM has been described in detail, some of the observed phenotypes associated with higher order mutants of these factors are not restricted to the primary root meristem (De Rybel et al., 2013; Ohashi-Ito et al., 2013a, 2013b). Moreover, there are some indications that TMO5 and LHW bHLH subclade members are expressed in aerial tissues as well (Ohashi-Ito et al., 2013a), consistently with the effect on SAM size observed in mutants and in the TMO5/LHW misexpression line **(Figure 1 A-E, K)**.

In order to first provide a detailed overview of the localisation of these factors in *Arabidopsis thaliana*, we generated promoter-nuclear triple GFP-GUS fusion constructs for all members and analysed their expression domains in the RAM, in the vascular cambium during root secondary growth and in the SAM. As previously reported (De Rybel et al., 2013), *TMO5* clade members show overlapping expression in the young xylem cells of the RAM (**Figure 3 A-D**). Similarly, during secondary growth in the root, *TMO5* clade members showed expression in developing and differentiating xylem cells. *T5L1* and *T5L3* were also detected in some cells of the vascular cambium (**Figure 3 E-H**). In the SAM region, only *TMO5* showed a specific provasculature-associated expression (**Figure 3I**). *T5L1* was not detected in the SAM (**Figure 3J**) but did show expression in a few cell files in the vascular tissue below the SAM (**Supplemental Figure S2A**). *T5L2* was found to be highly expressed in the L1 layer and at much lower levels in other cells of the SAM (**Figure 3K**). *T5L2* was also largely excluded from the centre of the meristem. *T5L3* was expressed broadly in the SAM both in the L1 layer and in the internal tissues, except for the central part of the SAM where it was completely absent (**Figure 3L**).

**Figure 3.**
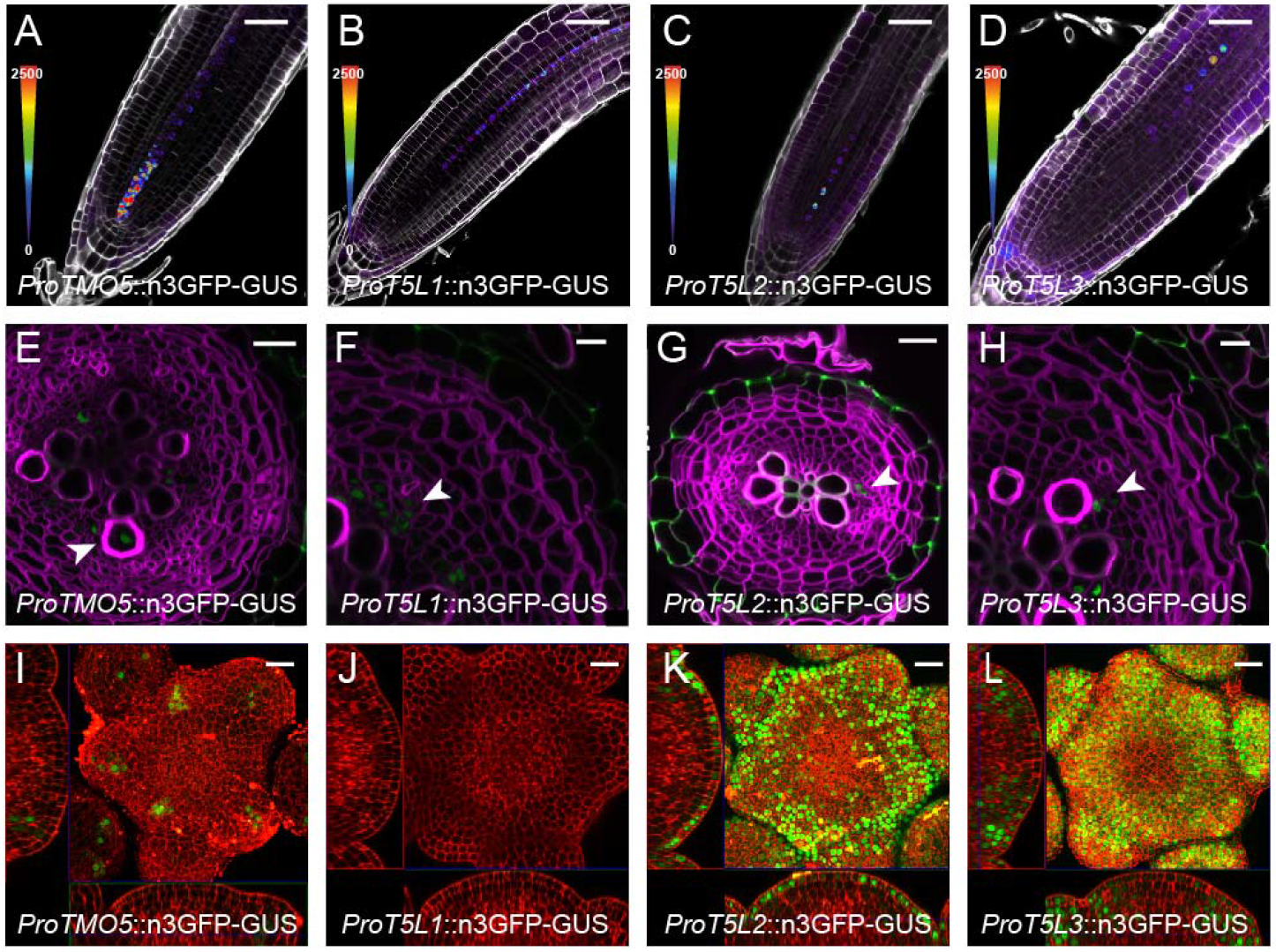
TMO5 clade members show specific expression patterns in the meristems. Promoter-reporter lines were used to analyse the expression pattern of *ProTMO5*, *ProT5L1*, *ProT5L2*, and *ProT5L3* in (A-D) longitudinal sections of 5-day-old root apical meristem; (E-H) cross sections of 20-day-old roots displaying secondary growth; (I-L) shoot apical meristems. Central squares in I, K and L represent maximum intensity projection. Scale bars: (A-D) 50 µm; (E, F, H) 10 µm; (G, I-L) 20 µm. Arrowheads indicate expression.

Compared to *TMO5* clade members in the RAM, members of the *LHW* clade showed a broader expression domain in vascular tissues (**Figure 4 A-D**), and in the case of *LHW* and *LL1*, this pattern was similar to previously published lines (**Figure 4 A, B**) (De Rybel et al., 2013). Also, during secondary growth, LHW, LL1 and LL3 clade members showed expression in xylem and cambium tissues, while LL2 was only detected in the xylem (**Figure 4 E-H**). A broad expression domain was observed in the SAM for both *LHW* and *LL1* (**Figure 4 I, J**), but *LHW* was absent specifically from the L2 layer while *LL1* was more specifically expressed in organ primordia in the peripheral zone. No expression was detected in the SAM for *LL2* and *LL3* (**Figure 4 K, L**) and, similarly to *T5L1*, *LL3* showed expression within the vasculature below the SAM (**Supplemental Figure S2B**). Taken together, our results show that the *TMO5* and *LHW* clade members are expressed in distinct meristematic regions, consistent with a general meristematic function for these factors throughout development. Additionally, some of these TFs show prominent expression only in one of the meristems, while being absent in others. These results thus argue for a general function of the TMO5-T5Ls/LHW-LLs factors in all the meristems, but with some expression specificity that could result in alternative TMO5-T5Ls/LHW-LLs combinations in the different meristems.

**Figure 4.**
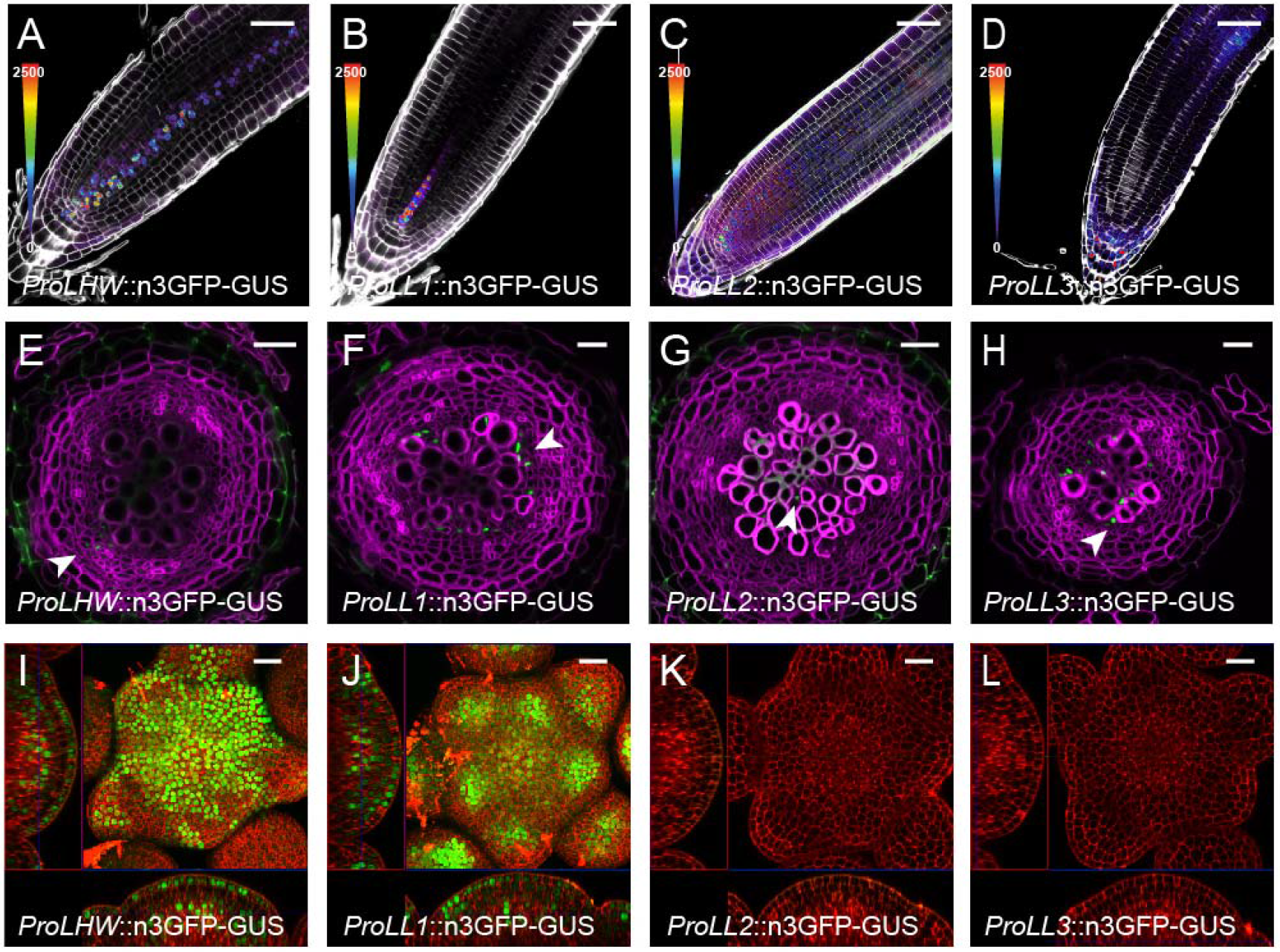
LHW clade members show overlapping expressions in the meristems. Promoter-reporter lines were used to analyse the expression pattern of *ProLHW*, *ProLL1*, *ProLL2*, and *ProLL3* in (A-D) longitudinal sections of 5-day-old root apical meristem; (E-H) cross sections of 20-day-old roots displaying secondary growth; and in (I-L) shoot apical meristems. Central squares in I and J represent maximum intensity projection. Scale bars: (A-D) 50 µm; (E-L) 20 µm. Arrowheads indicate expression.

### Single mutant analysis reveals functional specificity in TMO5 and LHW clade members

Despite the fact that TMO5 and LHW homologs have a clear redundant role in primary root vascular proliferation, the observed specificity in expression patterns in other meristems suggest that there might be some functional specificity amongst the clade members. Indeed, TMO5, T5L1 and LHW are reported to be the most prominent factors driving vascular proliferation in the primary root meristem, while the other members might be less important for this specific developmental process (De Rybel et al., 2013; Ohashi-Ito et al., 2014). To get a global understanding of the functional specificity among the TMO5 and LHW clade members throughout plant development, we next analysed single mutants for discernible phenotypes in the three meristems in comparison to WT plants. In the RAM, all single mutants with the exception of *t5l2* showed a significant reduction in cell files number (**Figure 5A, Supplemental Figure S3 A-I**). Still, a clear distinction within the subclades was observed in the relative contributions to this phenotype, with *tmo5*, *t5l1*, *t5l3* and *lhw* showing the strongest reduction in the number of vascular cell files. It is worth noting that *lhw* is the only single mutant with a monarch instead of the normal diarch vascular architecture (Ohashi-Ito and Bergmann, 2007), explaining the more pronounced phenotype. Similar observations were made in roots initiating secondary growth, with *tmo5, t5l1, t5l2, lhw, ll1* analysed mutants showing a significant reduction in cell file numbers, but the relative contributions of the factors were different. Indeed, *t5l1*, *t5l2* and *lhw* seem to be the major players in the establishment of secondary growth (**Figure 5B, Supplemental Figure S3 J-O**). Additionally, the SAM area was significantly larger in *t5l1* and *ll1* mutants as well as the number of meristem cells (**Figure 5C, Supplemental Figure S3 P-Y**). Cell size was not significantly affected in single mutants, confirming an effect on cell proliferation in the SAM (**Figure 5D**). Additional shoot phenotypes were found in the inflorescence stems of mutants. For example, *tmo5* and *t5l1* started initiating siliques before the last inflorescence branch (**Supplemental Figure S4A**) suggesting a deviation in lateral organ/structure identity determination in the SAM, whereas *t5l3* produced inflorescence branches incapable of upright growth (**Supplemental Figure S4B**). Finally, inflorescence growth of the *lhw* mutant was slower, and much more affected in *lhw ll1* double mutants (**Supplemental Figure S4A**). In summary, these results show that, while a general effect on cell proliferation is observed, the effect of single mutations in the members of *TMO5* and *LHW* subclades differs depending on the meristem considered, suggesting a level of functional and tissue specificity with an opposite trend in root and shoot.

**Figure 5.**
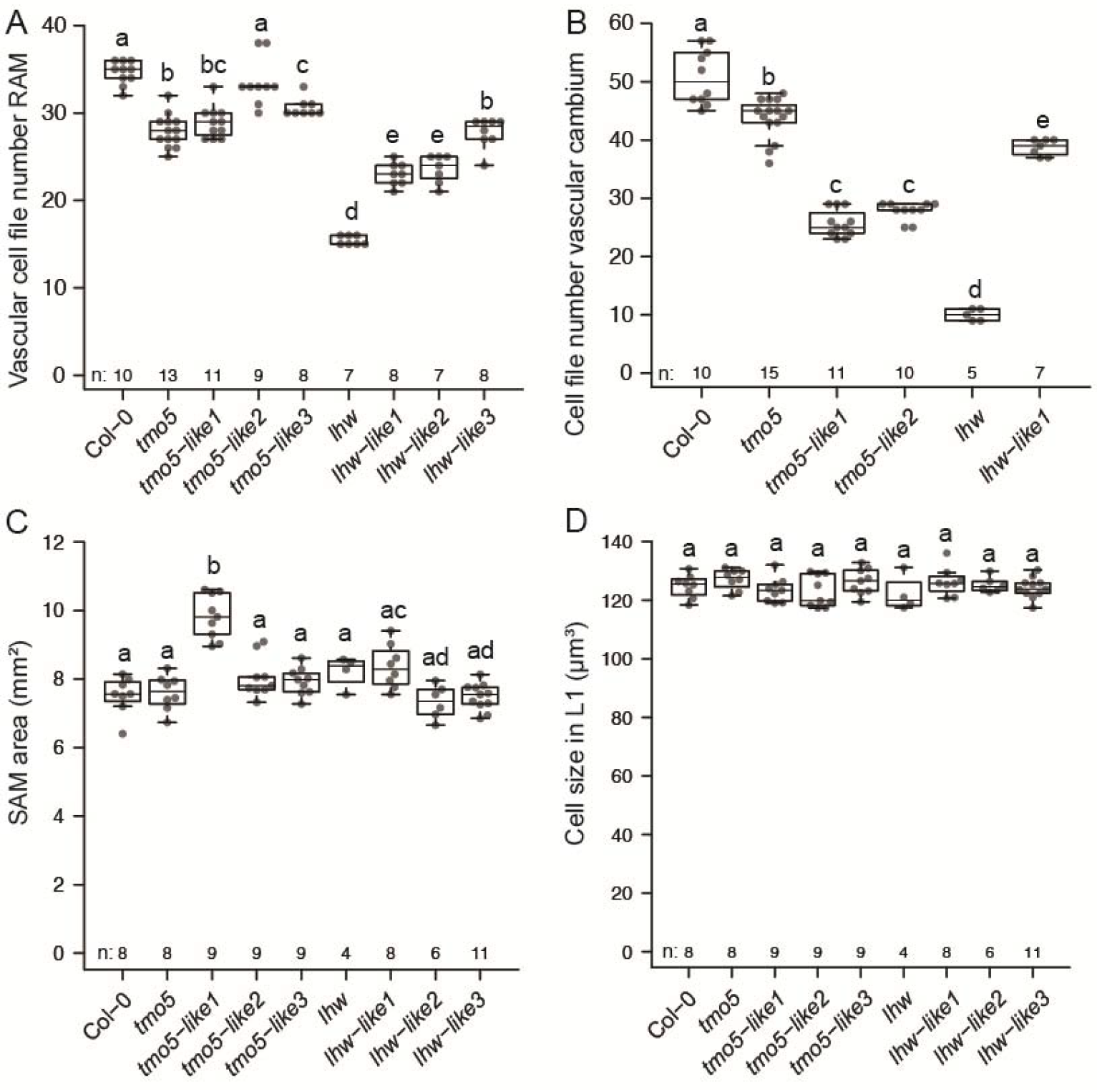
Single mutant analysis reveals functional specificity in TMO5 and LHW clade members. Quantification of Col-0, *tmo5*, *t5l1*, *t5l2*, *t5l3*, *lhw*, *ll1*, *ll2* and *ll3* for vascular cell files number of cross sections (A) of root apical meristems, (B) of root undergoing secondary growth, (C) measurement of shoot apical meristem area and (D) cell size in L1 layer. Asterisks indicate endodermis. Lowercase letters in charts indicate significantly different groups as determined using a one-way ANOVA with post-hoc Tukey HSD testing (p ≤ 0.05).

### Variations in bHLH heterodimer complexes show distinct phenotypic outputs

Although the observed tissue- and organ-specific expression of the TMO5 and LHW clade members could account for the phenotypic differences in the single mutants, there was no perfect correlation, suggesting that there might be other layers of functional regulation. Given that TMO5 and LHW clade members form obligate heterodimer complexes (De Rybel et al., 2013; Ohashi-Ito et al., 2014), we next questioned if such functional specificity could be caused by the particular heterodimer complex that is formed. Although TMO5 and LHW interaction partners can likely all interact with each other (De Rybel et al., 2013; Ohashi-Ito et al., 2014), this indeed does not mean that these combinations would lead to a functional bHLH complex. We thus combined several individual overexpression lines by crossing TMO5-clade members with LHW-clade members and analysed the effect in the RAM and the SAM (**Figure 6**). Compared to the wild type, different combinations resulted in a quantitative difference in the number of root vascular cell files with TMO5/LHW as the most potent combination and T5L2/LHW not showing any significant difference (**Figure 6 A-E and K**). Strong phenotypical effects were observed in the shoot as previously reported for TMO5/LHW (Vera-Sirera et al., 2015), including reduced stem height, curly, hyponastic, or jagged leaves (**Supplemental Figure S5**). In the SAM, our analysis suggests that the resulting phenotypes and their severity show a tendency to be dependent on the combination of TMO5- and LHW-clade members as in the root but the high variability of the parameters allows firm conclusions only for TMO5/LL2 with a reduced SAM area (**Figure 6 F-J and L**) and SAM cell number **(Supplemental Figure S6)**, and TMO5/LHW with a reduced cell size **(Figure 1M and 6M).**

**Figure 6.**
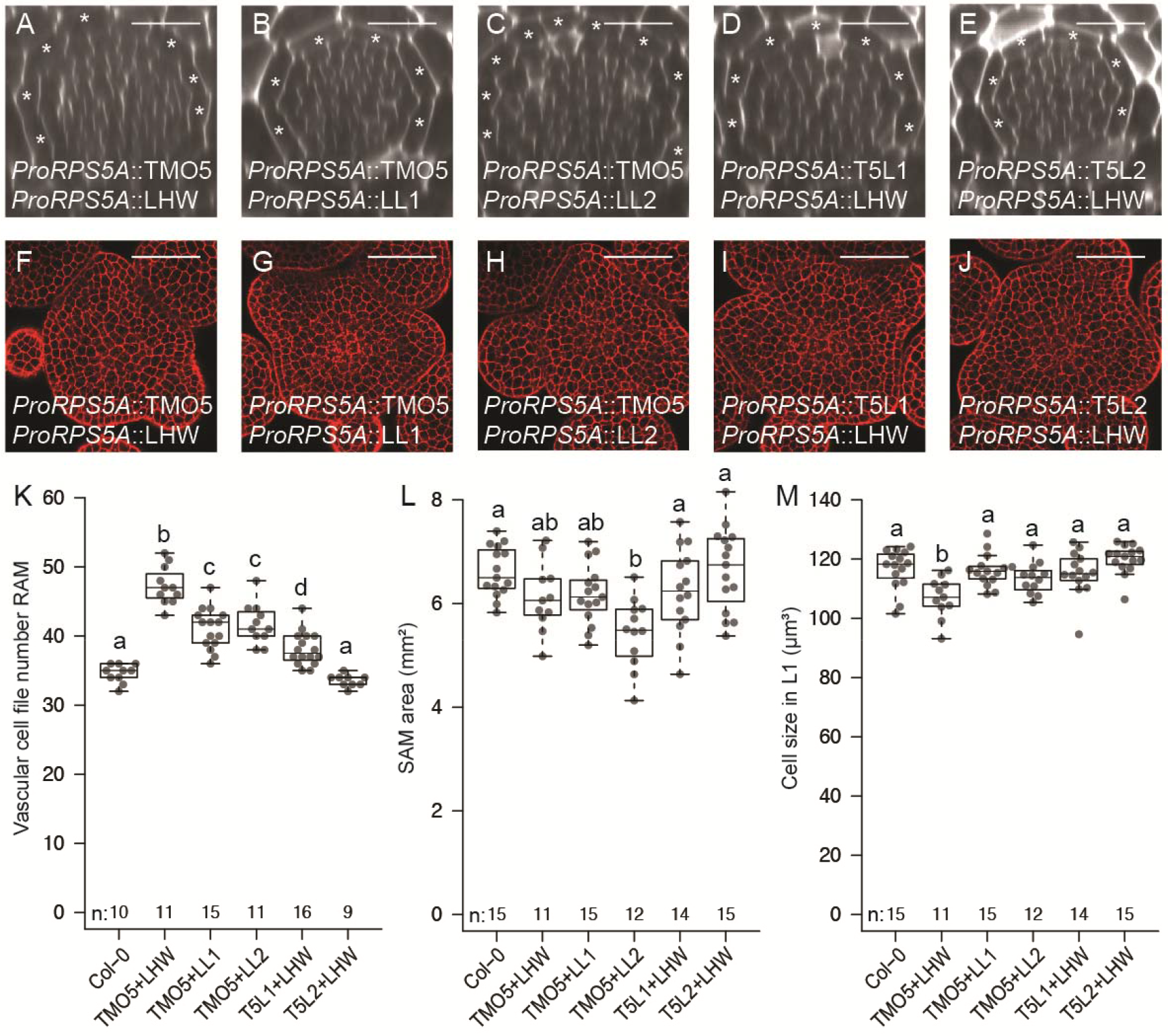
Variations in bHLH heterodimers show distinct phenotypic outputs. Ortho-views of z-stack confocal microscopy images of *ProRPS5A*::TMO5 x *ProRPS5A*::LHW; *ProRPS5A*::TMO5 x *ProRPS5A*::LL1; *ProRPS5A*::TMO5 x *ProRPS5A*::LL2; *ProRPS5A*::T5L1 x *ProRPS5A*::LHW and *ProRPS5A*::T5L2 x *ProRPS5A*::LHW of (A-E) RAM of 5-day-old seedlings, and (F-J) shoot apical meristems. (K) Quantification of vascular cell files number of RAM, (L) measurement of shoot apical meristem area and (M) cell size in L1 layer. Lowercase letters in charts indicate significantly different groups as determined using a one-way ANOVA with post-hoc Tukey HSD testing (p ≤ 0.05). Scale bars: 20 µm.

In summary, these experiments show that even when TMO5 and LHW clade members are ectopically expressed together, they do not always lead to the same phenotype. Thus, TMO5- and LHW-clade heterodimer complex activity is not solely determined via the observed differential expression domains but also likely because of variations in the complexes or in the activity of the complexes which are being formed.

### Variations in bHLH heterodimer complexes affect target gene specificity

TMO5/LHW complexes induce vascular cell proliferation in the root apical meristem via induction of cytokinin biosynthesis through direct binding to the promoter regions of *LOG3* and *LOG4* (De Rybel et al., 2014; Ohashi-Ito et al., 2014; Smet et al., 2019). These enzymes catalyse the final step of cytokinin biosynthesis (Kurakawa et al., 2007; Kuroha et al., 2009). We therefore set to explore the functional differences in gene regulatory potential between the different heterodimers that can be formed. To achieve this, we implemented a mid-throughput protoplast-based quantitative gene expression system that enables covering all possible combinations, obtaining quantitative data and reducing interferences from other factors and tissue context. Protoplasts derived from Arabidopsis shoots were transiently transformed with constructs comprising each combination of the TFs under the control of a 35S constitutive promoter. The promoters of the tested genes, namely *LOG1*, *LOG3*, *LOG4*, *LOG5* and *LOG7*, were cloned upstream the firefly luciferase gene, that served as a readout. A construct coding for constitutively expressed Renilla luciferase was included as a normalization element. We first assayed the effect of overexpression of single TFs on the expression of each of the *LOG* genes (**Supplemental Figure S7A**). Only LHW and LL1 were able to induce *LOG3* and *LOG4* expression to moderate levels, whereas all other TFs did not. These results are overall consistent with the idea that TMO5 and LHW clade members act as obligate heterodimers (De Rybel et al., 2013; Ohashi-Ito et al., 2014). It also suggests that LHW and LL1 can activate a basal level of transcription by themselves. Overexpression of any combination of two transcription factors from the TMO5 and LHW clades was able to induce expression of the direct target genes *LOG3* and *LOG4*, but there was a clear quantitative difference with the highest induction values for all TMO5-clade combinations with LL1, and the lowest in combinations with LL2 (**Figure 7**). Besides the clear quantitative effect for *LOG3* and *LOG4* expression values, other *LOG* genes analysed were not induced (**Figure 7**), suggesting a specific regulation of LOG3 and LOG4 by the TMO5- and LHW-clade members or that other regulatory factors not present in protoplasts might be needed for their induction. Moreover, these experiments show that rather the LHW-clade members play a key role in defining the strength of the transcriptional activation in this simplified system, not the TMO5 clade.

**Figure 7.**
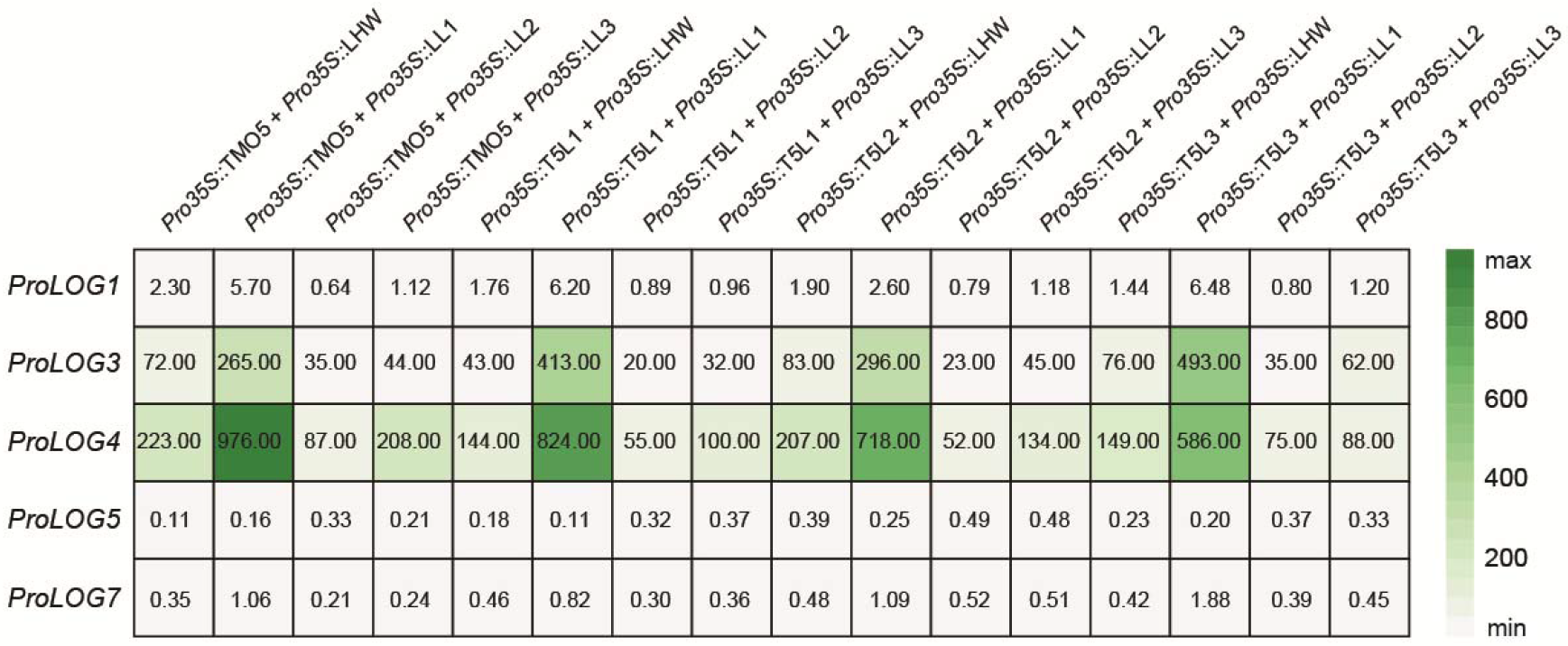
Different combinations in bHLH heterodimer complexes affect target gene expression. Heat map shows relative changes of gene expression from *LOG* promoters after overexpression of combinations of TMO5 and LHW clade members’ pairs in quantitative gene expression assays in Arabidopsis protoplasts. Values represent FLuc/RLuc ratios, n = 4-6.

To further validate these results *in planta*, we inspected the *ProLOG3::n3GFP* and *ProLOG4::n3GFP* reporter lines in *lhw* and *tmo5 t5l1* mutant backgrounds (**Supplemental Figure S7 B-G**). In a control line, *ProLOG3::n3GFP* is expressed in a diarch configuration in primordia (**Supplemental Figure S7B**; white arrows), similarly to a diarch expression in root protoxylem cells (De Rybel et al., 2014). The signal of *ProLOG3::n3GFP* was still present in the *tmo5 t5l1* double mutant (**Supplemental Figure S7C**) despite a loss of activity of two TMO5 clade members affecting SAM phenotype (**Figure 1 C, L; Supplemental Figure S4**). However, the expression of *ProLOG3::n3GFP* in *tmo5 t5l1* was detected only in one cell axis compared to the control in an inspected SAM area (**Supplemental Figure S7C**; yellow arrows) suggesting impaired vascular formation similar to the effects in the RAM (Ohashi-Ito and Bergmann, 2007; De Rybel et al., 2013). The expression of *ProLOG4::n3GFP* in the L1 layer (**Supplemental Figure S7E**) decreased in a *tmo5 t5l1* double mutant but the overall pattern remained the same (**Supplemental Figure S7F**). In contrast, the signal of both *ProLOG3::n3GFP* and *ProLOG4::n3GFP* was completely missing in a *lhw* single mutant (**Supplemental Figure S7 D, G)**. Overall, these findings validate the results of the protoplast assays showing that LHW activates *LOG3* and *LOG4* transcription together with TMO5 clade members, and that the LHW-clade members are essential for *LOG3* and *LOG4* expression.

## CONCLUSIONS AND DISCUSSION

Current knowledge on the regulatory complexes governing cell proliferation suggests that plants have evolved specific networks for each of the meristem regions. Although most known examples are comparable since they are based on a peptide-receptor interaction pair, these are unique for one specific meristem context: CLV3-CLV1 in the SAM (Gaillochet et al., 2015), CLE41-PXY in the vascular cambium (Fisher et al, 2007; Suer et al., 2011; Etchells et al., 2013), and CLE40-ACR4 in the RAM (Stahl et al., 2013; Berckmans et al., 2020). One can however question if this is accurately reflecting an evolutionary reality where each meristem region has independently evolved a dedicated regulatory network, or whether we are simply yet to uncover common regulators in these distinct regions governing cell proliferation. Our results suggest that the TMO5/LHW and T5Ls/LLs bHLH heterodimer complexes act as general regulators of cell proliferation expressed and active in all plant meristems. In this case, heterodimer variations by combinations of TMO5 and LHW subclade members resulting primarily from expression in specific domains within different meristems (and not from differential expression between meristems) provide the required specificity to adapt responses to a given developmental context. Although all homologs show overlapping expression domains in the RAM, there is more variation in expression domains in the SAM and vascular cambium areas (**Figure 3 and 4**). This highlights the fact that conservation of expression patterns in one developmental system is not necessarily copied to other contexts, further complicating extrapolation of functional studies performed in one organ to another. The obligate heterodimer nature of this interaction is likely to be of key importance here, as homodimers present in e.g. single misexpression lines (De Rybel et al., 2013) do not give the strong phenotypical effect as when both partners are overexpressed. Moreover, heterodimers are required for an efficient regulation of the well characterized target genes *LOG3* and *LOG4* (**Figure 7; Supplemental Figure S7**). Additionally, higher order mutants of each subclade, such as *tmo5 t5l1 t5l2 t5l3* and *lhw ll1* mutants, yield the same phenotype (De Rybel et al., 2013). Our results however suggest that the activity of TMO5/LHW heterodimers is also further determined by specificity of the promoter regions, restricting expression of a few TMO5/LHW homologs to one or more meristems. Some TMO5/LHW combinations are thus also likely to act specifically in a given meristem.

An intriguing question emerging from our results is whether the bHLH heterodimer complexes are unique in their capacity to act as more general regulators that can be used in different developmental contexts; or whether this is a more general theme for most TFs which has simply not been uncovered so far. On one hand, one could argue for bHLH factors being unique as there are other examples of a same set of bHLH factors acting in multiple contexts. For example, the formation of trichomes and root hairs respectively depends on the bHLH proteins GL3 or EGL3, which interact with the MYB proteins WER or GL1 thus forming active transcriptional complexes used in these two developmental contexts (Bernhardt et al., 2003; Zhang et al., 2003; Zhao et al., 2008). On the other hand, as a general property, several TF families can form within-family heterodimers and TFs from a given family can interact with many other TFs from other families (Trigg et al., 2017). Thus, variations of heterodimer formation of TFs from other families than bHLH could occur in different meristems through differential expression and could be regulating meristems similarly to the TMO5/LHW heterodimer. Although it might thus be evolutionary efficient for organisms to use TFs dedicated to a given tissue, combinatorial TF interactions among different families is an alternative to achieve the same level of specificity needed in each developmental context through regulation of expression patterns within and not between tissues. It is thus likely that we are yet to uncover additional functions of known TFs in other developmental contexts which could emerge by interactions with other partners.

Although our results suggest that TMO5/LHW heterodimers act as a common regulator controlling cell proliferation, the exact downstream mechanism might not be conserved in each of the meristem contexts. *LOG* genes are induced by all tested heterodimer complexes in a protoplast system and complexes containing LHW, and LL1 are required for their expression both in the RAM and SAM (De Rybel et al., 2014) (**Supplemental Figure S7**). This suggest that regulation of cytokinin biosynthesis could be a conserved mechanism between the different meristems. However, opposite responses on cell proliferation are observed upon misexpression in the SAM and RAM regions, which could be due to the differences in DNA binding specificity of the different heterodimers in each developmental context. It is very likely that a different set of target genes will be activated by specific heterodimer complexes leading to further functional diversification. Some indications can be found in literature as T5L1/LHW misexpression was shown to have only partly overlapping target genes compared to TMO5/LHW misexpression (De Rybel et al., 2014; Ohashi-Ito et al., 2014). However, these gene lists have been obtained using a very different experimental set-up, precluding a direct comparison. As such, additional work using a genome-wide analysis of target genes for the different heterodimer complexes would be required to evaluate the precise downstream target gene sets activated by these complexes. In summary, we show that a common bHLH heterodimer complex module controls cell proliferation in distinct plant meristems in *Arabidopsis thaliana* through heterodimer diversification leading to target gene specification.

## METHODS

### Plant material

Unless otherwise mentioned, all plant material used was *Arabidopsis thaliana*, ecotype Columbia-0. Some transgenic and mutant lines have been described previously: *ProLOG3::n3GFP* (De Rybel et al., 2014), *ProLOG4::n3GFP* (De Rybel et al., 2014), *ProRPS5A::TMO5-GR* (De Rybel et al., 2013), *ProRPS5A::TMO5-GR* x *ProRPS5A::LHW-GR* (Smet et al., 2019). *ProRPS5A* overexpression lines (De Rybel et al., 2013) were used to generate the crosses: F1 seeds were used for RAM analysis, each seedling was genotyped to confirm the presence of the constructs. F2 seeds from genotyped plants were used for the SAM analysis. The n3GFP-GUS reporter lines were generated by MultiSite Gateway cloning (Karimi et al., 2007) into the pMK7S*NFm14GW,0 destination vector. All constructs were transformed into the *Arabidopsis thaliana* Col-0 background.

For phenotype analyses, we used the following mutant lines: *tmo5* (GK-143E03) (De Rybel et al., 2013), *t5l1* (RIKEN_12-4602-1) (De Rybel et al., 2013), *t5l2* (GK-824H07), *t5l3* (SALK_109295) (De Rybel et al., 2013), *lhw* (SALK_023629) (De Rybel et al., 2013), *ll1* (SALK_126132) (Ohashi-Ito et al., 2013a), *ll2* (GK-523B12), *ll3* (GK-262H03). Gene specific primers for genotyping were designed and are listed in **Supplemental Table S2**, as are insertion-specific primers. For expression analysis *in planta*, *ProLOG3::n3GFP* and *ProLOG4::n3GFP* lines were crossed into *lhw* or *tmo5 t5l1* mutant backgrounds. Homozygous plants were selected by PCR or antibiotic resistance as follows: final concentrations in cultivation medium of 25 mg/l kanamycin (Duchefa), 20 mg/l hygromycin (Duchefa), 10 mg/l sulfadiazine (Sigma-Aldrich). *ProLOG4::n3GFP* in *lhw* background was published previously (De Rybel et al., 2014).

### Cultivation conditions

For root analysis, seeds were sterilized using a solution of 25% bleach and 75% ethanol. After 4 days of stratification at 4°C, plants were grown in half strength Murashige and Skoog medium (Duchefa) (Murashige and Skoog, 1962) without sugar and 0.8% Plant agar under continuous light conditions at 22°C. 10 µM dexamethasone (DEX) was used for induction of expression. For lateral meristem root analysis, plants were grown in half strength Murashige and Skoog medium (Duchefa) and 0.8% plant agar under continuous light conditions for 19-20 days at 22°C. For shoot analysis, plants were cultivated in soil under long-day conditions (16 hours light/8 hours dark) in growth chambers maintained at 21-22°C, with a light intensity of approximately 150 μmol m^-1^ s^-1^ and 40-60% relative humidity.

### Shoot apical meristem dissection

Shoot apical meristems from inflorescence stems between 0.5 and 1.5 cm long were dissected and cultured *in vitro* for 3 hours in a cultivation chamber as described previously (Brunoud et al., 2020). The meristems were stained with a water solution of 100 µg/ml propidium iodide (Sigma-Aldrich) for 5 minutes, then washed with water and used for microscopy.

### Histochemical and histological procedures

For anatomical sections, 10-days-old roots were fixed overnight in 1% glutaraldehyde and 4% paraformaldehyde in 50 mM phosphate buffer, pH 7. Samples were dehydrated and embedded in Technovit 7100 resin (Heraeus Kulzer) according to the manufacturer’s protocol. For proper orientation of the samples, we used a two-step embedding methodology, with a pre-embedding step to facilitate orientation in 0.5 ml Eppendorf tubes (De Smet et al., 2004). Sections of 4 μm of root, taken 0.5 cm below junction between the root and the hypocotyl were cut with a Richert Jung microtome 2040, dried on Superfrost® plus microscopic slides (Menzel-Gläser), counterstained for cell walls with 0.05% ruthenium red for 5 minutes and rinsed in water. After drying, the sections were mounted in DPX mounting medium (Sigma-Aldrich) and covered with cover slips. Images were taken with an Olympus BX53 DIC microscope. mPS-PI staining was performed as described previously (Truernit and Haseloff, 2008). Briefly, the seedlings were fixed in 50% methanol and 10% acetic acid at 4°C for at least 12 hours. Samples were then rinsed with water and incubated in 1% periodic acid (Sigma-Aldrich) for 40 minutes at room temperature (22°C). After another water rinse, seedlings were incubated with Schiff’s reagent (100 nM sodium metabisulphite, 0.15N 37% HCl) with fresh propidium iodide (100 µg/ml) until visibly stained. To visualize, seedlings were transferred onto microscope slides in chloral hydrate solution. Quantification of vascular cell file numbers (cells within but excluding the pericycle) were performed using ImageJ software (https://imagej.nih.gov/ij/). Root apical meristems of n3GFP-GUS seedlings were stained with 0.1% Calcofluor White in ClearSee solution to visualize the cell wall (Ursache et al., 2018). To visualize GFP during secondary growth, a modified ClearSee protocol (Ursache et al., 2018; Ben-Targem et al., 2021) was employed. The most upper part of the root (0.5 cm below the hypocotyl root junction) was fixed with 4% PFA (Paraformaldehyde: Sigma, P6148) and 0.01% Triton in 1x PBS for 1 hour under vacuum and embedded in 5% agarose blocks, then sections of 70-80 µm were obtained using a Vibratome (Leica VT-1000) and collected in water. Water was quickly replaced with ClearSee solution (10% xylitol 15% sodium deoxycholate, 25% urea) (Kurihara et al., 2015) and sections were kept in ClearSee for 24 hours at room temperature and then stored at 4°C. Prior imaging, sections were stained with 0.05% Direct Red 23 (Sigma 212490) in ClearSee for 30 minutes, washed 3 times in ClearSee and mounted in ClearSee on a slide. Direct Red 23 stained the cell wall, and it was used to visualized cell outlay.

### Microscopy

Confocal microscopy of shoot apical meristems was carried out using an upright Zeiss Axio Imager 2 equipped with a LSM700 confocal unit and 40x/1.0 DIC M27 water-dip objective. GFP was excited at 488 nm and detected at 490-530 nm; PI was excited at 555 nm and detected at 570-630 nm. Confocal microscopy of n3GFP-GUS root apical meristems was performed on a Leica SP8 using a 63x water-immersion objective. Calcofluor White and GFP were excited at 405 nm and 488 nm and visualized at 425-475 nm and 500-550 nm, respectively. mPS-PI-stained roots were imaged at an excitation of 514 nm and emission of 600-650 nm. Confocal microscopy of root sections undergoing secondary growth was performed on a Zeiss LSM880 using a 20x dry objective and digital zoom. GFP and Direct Red 23 were excited at 488 nm and 561 nm and visualized at 490-544 nm and 580-642 nm, respectively. DIC microscopy of embedded samples was done using an Olympus BX53 microscope equipped with 10x, 20x and 40x air objectives.

### cDNA synthesis

Total RNA was prepared from 100 mg of 11-day-old seedlings with the NucleoSpin RNA Plant Kit (Macherey-Nagel) according to the manufacturer’s instructions. cDNA was synthesized from 500 ng of total RNA using the SuperScript™ III Reverse Transcriptase Kit (Invitrogen).

### Plasmid construction

DNA fragments were released by restriction from existing plasmids or amplified by PCR using primers synthesized by Sigma-Aldrich or Eurofins. The PCR reactions were performed using Q5 High-Fidelity DNA Polymerase (New England Biolabs). Gel extractions were performed using NucleoSpin Gel and PCR Clean-up Kits (Macherey-Nagel). Vectors were assembled via AQUA cloning technology (Beyer et al., 2015) and transformed into chemically competent *E. coli* strain 10-beta (New England Biolabs) or TOP10 (Invitrogen). Plasmid purifications were performed utilising Wizard Plus SV Minipreps DNA Purification Systems (Promega). New plasmids were tested by restriction enzyme digests and sequencing (Eurofins/GATC or Microsynth). All restriction enzymes were purchased from New England Biolabs. For testing promoters, the plasmid pMP010 was constructed as follows: the firefly luciferase gene (*FLuc*) was amplified by PCR from the pMZ836 plasmid (Müller et al., 2014) using the oligonucleotides oMP025 and oMP029. The product was assembled via AQUA cloning into pGEN16 (Samodelov et al., 2016) digested with SacII/XhoI. Promoter sequences of LOG genes upstream from the ATG were amplified from genomic DNA extracted from 7-day-old seedlings using primers as follows: ProLOG1 (oMP036 and oMP037, 3138 bp), ProLOG3 (oMP040 and oMP041, 3564 bp), ProLOG4 (oMP022 and oMP023, 3999 bp), ProLOG5 (oMP042 and oMP043, 3024 bp), ProLOG7 (oMP026 and oMP027, 3187 bp). The products were inserted via AQUA cloning into pMP010 digested with SacII/AgeI. For preparing vectors harbouring cDNA of transcription factors, the plasmid pMP011 was constructed as follows: the nucleotide sequence of the HA tag (YPYDVPDYA) was amplified by PCR using the oligonucleotides oMP088 and oMP089. The product was assembled via AQUA cloning into pGEN16 digested with AgeI/XhoI. Nucleotide sequences of transcription factors were amplified from cDNA prepared previously using primers as follows: cTMO5 (oMP107 and oMP108), cT5L1 (oMP121 and oMP122), cT5L2 (oMP125 and oMP126), cT5L3 (oMP123 and oMP124), cLHW (oMP109 and oMP110), cLL1 (oMP131 and oMP132), cLL2 (oMP129 and oMP130), cLL3 (oMP127 and oMP128). PCR products were fused via AQUA cloning into pMP011 digested with AfeI/BstZ17I. All primers and plasmids used in this study are listed in **Supplementary Table S2** and **Supplementary Table S3**, respectively.

### Luciferase protoplast assay

Protoplasts were isolated from shoots of 2- to 3-week-old *Arabidopsis thaliana* plants. Floatation was employed for isolation, and plasmids were transformed using a polyethylene-glycol-mediated approach as described previously (Ochoa-Fernandez et al., 2020). Plasmids were prepared with a Wizard® Plus Midipreps DNA Purification System (Macherey-Nagel). Protoplasts were co-transformed with mixtures of the appropriate plasmids, 30 μg DNA in total. The transformed protoplasts were cultivated for 18–20 hours at 19–22°C in the dark. After incubation, protoplasts were divided into aliquots of sufficient volume to measure six technical replicates for each sample. Firefly (FLUC) and Renilla luciferase (RLuc, in GB0109, (Sarrion-Perdigones et al., 2013)) activities were simultaneously quantified in intact protoplasts as described (Ochoa-Fernandez et al., 2016). Substrates for both luciferases were added directly before measurement: D-luciferin (Biosynth AG) for FLuc, Coelenterazine (Carl Roth) for RLuc. Chemiluminescence measurements were performed using a Berthold Centro XS3 LB 960 microplate luminometer (Berthold Technologies, Bad Wildbad, Germany) and a BertholdTriStar2 S LB 942 multimode plate reader (Berthold Technologies, Bad Wildbad, Germany). The FLuc/RLuc ratio was determined (n = 4–6) and showed in tables. Constitutively expressed RLuc served as an internal normalization element to obtain ratiometric data.

### Quantitative analysis of shoot apical meristem

Images of shoot apical meristems stained with propidium iodide were segmented using an auto seeded 3D watershed algorithm derived from the MARS pipeline (Fernandez et al., 2010) in which the parameters were manually tuned for each sample. In the resulting segmented images, cells belonging to the L1, L2 and L3 layers were automatically identified. To do so, a triangle mesh representing the tissue surface was computed using the segmented image, and L1 cells were selected as those adjacent to the background region and closest to the vertices of the surface mesh. L2 and L3 cells were selected recursively by adjacency to cells belonging to the previous layer.

Finally, “meristematic cells” (cells belonging to the central zone, the peripheral zone and to organ initials) were distinguished from cells of organ primordia and boundaries using the surface curvature. Principal curvatures were estimated on the surface mesh based on the vertex normal vectors (Theisel et al., 2004), and a central meristematic region was identified by thresholding the minimum principal curvature value and performing morphological operations. The retained threshold value was −0.005μm^−1^. The resulting binary property was projected on the closest L1 cells to identify L1 meristematic cells, and then propagated to L2 and L3 cells by adjacency with a triangle of meristematic cells at the previous layer. The results were obtained by filtering out non-meristematic cells and pooling the cell measures by cell layer.

### Statistical analysis

All statistical analysis plots were generated using the PlotsOfData webtool at standard settings (https://huygens.science.uva.nl/PlotsOfData/). In all boxplots, boxes represent the 1st and 3rd quartiles, and the centre line represents the median. The lowercase letters associated with the boxplots indicate significantly different groups as determined by one-way analysis of variance (ANOVA) with post-hoc Tukey HSD testing (P<0.001).

### Accession Numbers

The sequence data of genes described this article can be found in The Arabidopsis Information Resource (https://www.arabidopsis.org/) or GenBank (http://www.ncbi.nlm.nih.gov/genbank/) databases under the following accession numbers: *AT3G25710* for *TMO5/bHLH32*, *AT1G68810* for *T5L1/bHLH30/ABS5*, *AT3G56770* for *T5L2/bHLH107*, *AT2G41130* for *T5L3/bHLH106/STC8*, *AT2G27230* for *LHW/bHLH156*, *AT1G64625* for *LL1/LHL3/bHLH157*, *AT2G31280* for *LL2/LHL2/bHLH155*, *AT1G06150* for *LL3/LHL1/EMB1444*, *AT2G28305* for *LOG1*, *AT2G37210* for *LOG3*, *AT3G53450* for *LOG4*, *AT4G35190* for *LOG5*, and *AT5G06300* for *LOG7*.

## ACKNOWLEDGMENTS

We would like to acknowledge the Core Facility CELLIM (supported by MEYS CR, project LM2018129 Czech-BioImaging) and Plant Sciences Core Facility of CEITEC Masaryk University as well as SFR Biosciences (UMS3444/CNRS, US8/Inserm, ENS de Lyon, UCBL) PLATIM microscopy facility for their valuable technical support. This project has received funding from the European Structural and Investment Funds, Operational Programme Research, Development and Education (project „MSCAfellow@MUNI“, number CZ.02.2.69/0.0/0.0/17_050/0008496) to M.P, the European Research Council grant (ERC; StG TORPEDO; 714055) to E.M., the Ghent University Special Research Fund (BOF20/GOA/012) to M.M., and the DFG ( RA2950/1-2 and INST 37/965-1 FUGG) to L.R., the German Research Foundation (DFG) under Germanýs Excellence Strategy (CEPLAS - EXC-1028 project no. 194465578 and EXC-2048/1 – Project no. 390686111), the iGRAD Plant (IRTG 1525) and NEXTPlant (Project ID 391465903/GRK 2466) to MDZ.

## AUTHOR CONTRIBUTIONS

EM, MP, BDR and TV designed the research. EM performed the experiments related to RAM and vascular cambia with the help of MM and JN. LR and DR analysed root secondary growth expression pattern. MP designed and performed the experiments related to SAM, shoot phenotype and LUC assays with the help of GC, CG, JA and MDZ. EM and MP prepared figures and manuscript draft. EM, MP, BDR and TV wrote the manuscript with input from all authors.

---

**Supplemental Figure S1.**
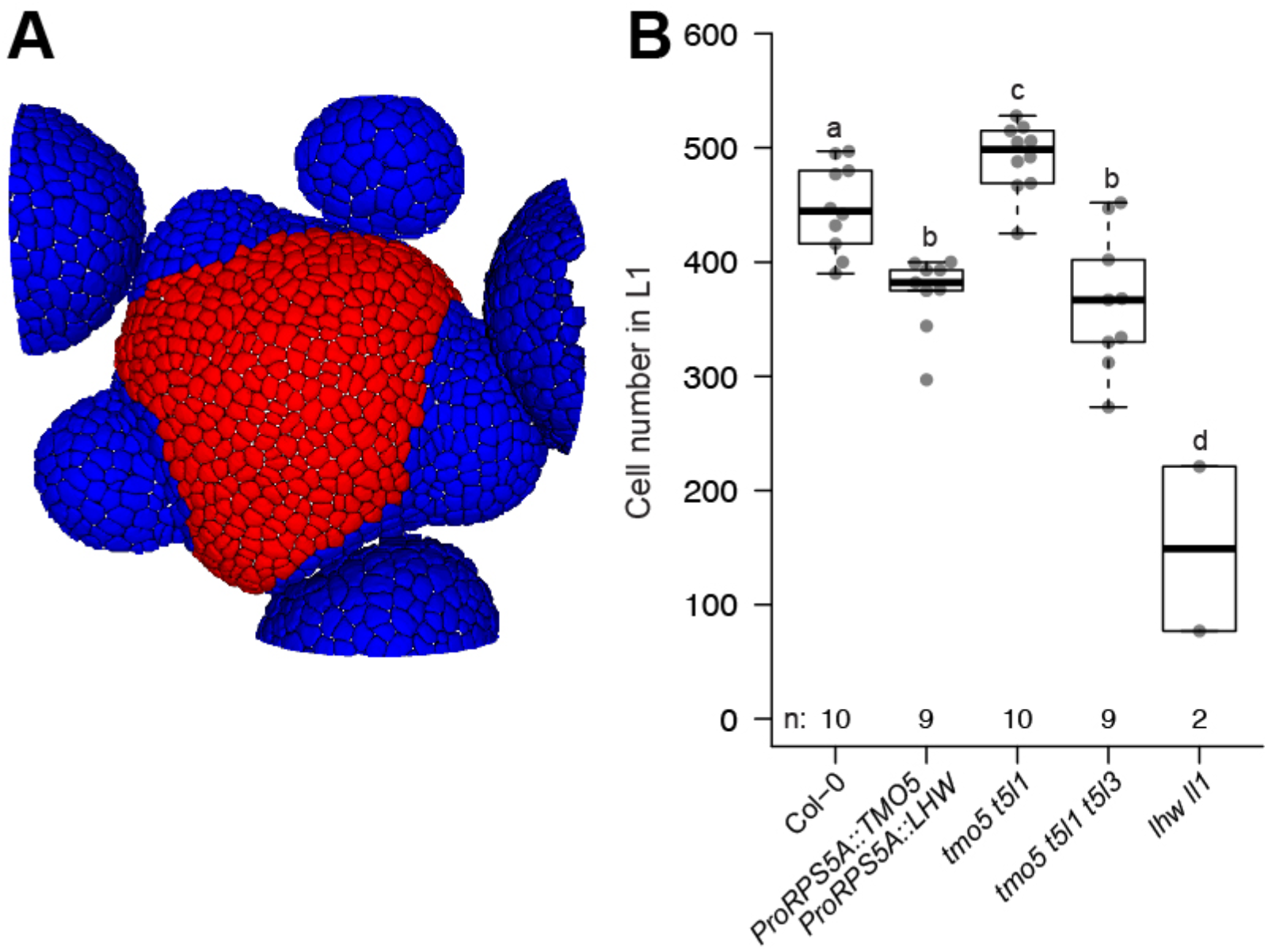
Quantification of cell number in TMO5/LHW overexpression line and multiple mutants. (A) An example of a SAM surface used for quantification of SAM parameters. Cells for determination of SAM area are depicted in red. (B) Cells were counted in L1 layer of shoot apical meristems. Lowercase letters indicate significantly different groups as determined using a one-way ANOVA with post-hoc Tukey HSD testing (p ≤ 0.05).

**Supplemental Figure S2.**
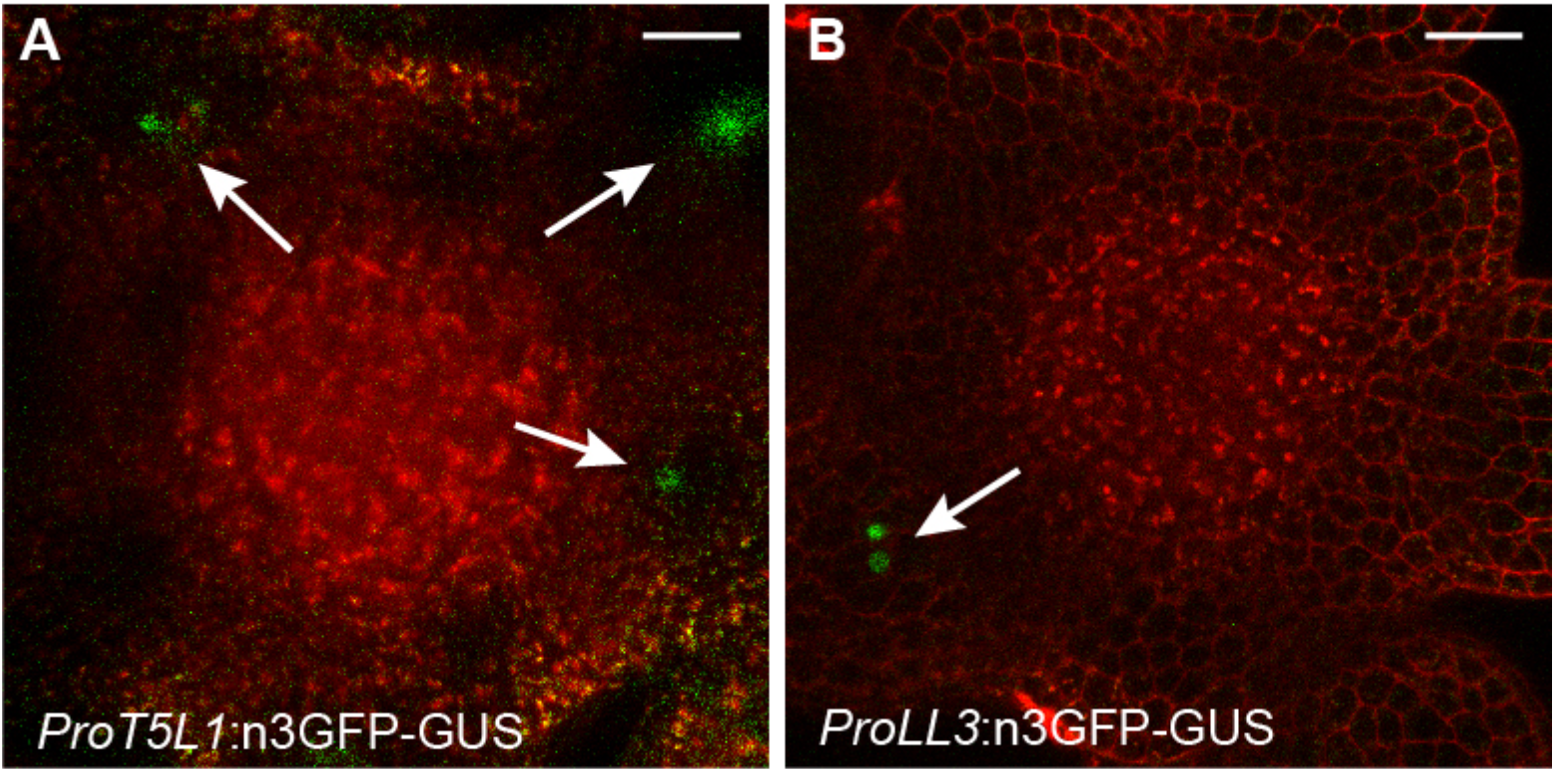
Expression pattern of *ProT5L1* and *ProLL3* in shoot apical meristem. Confocal microscopy images of shoot apical meristems of (A) *ProT5L1* and (B) *ProLL3* GFP-GUS reporter lines. Arrows indicate signal in deeper plant tissues. Scale bars: 20 µm.

**Supplemental Figure S3.**
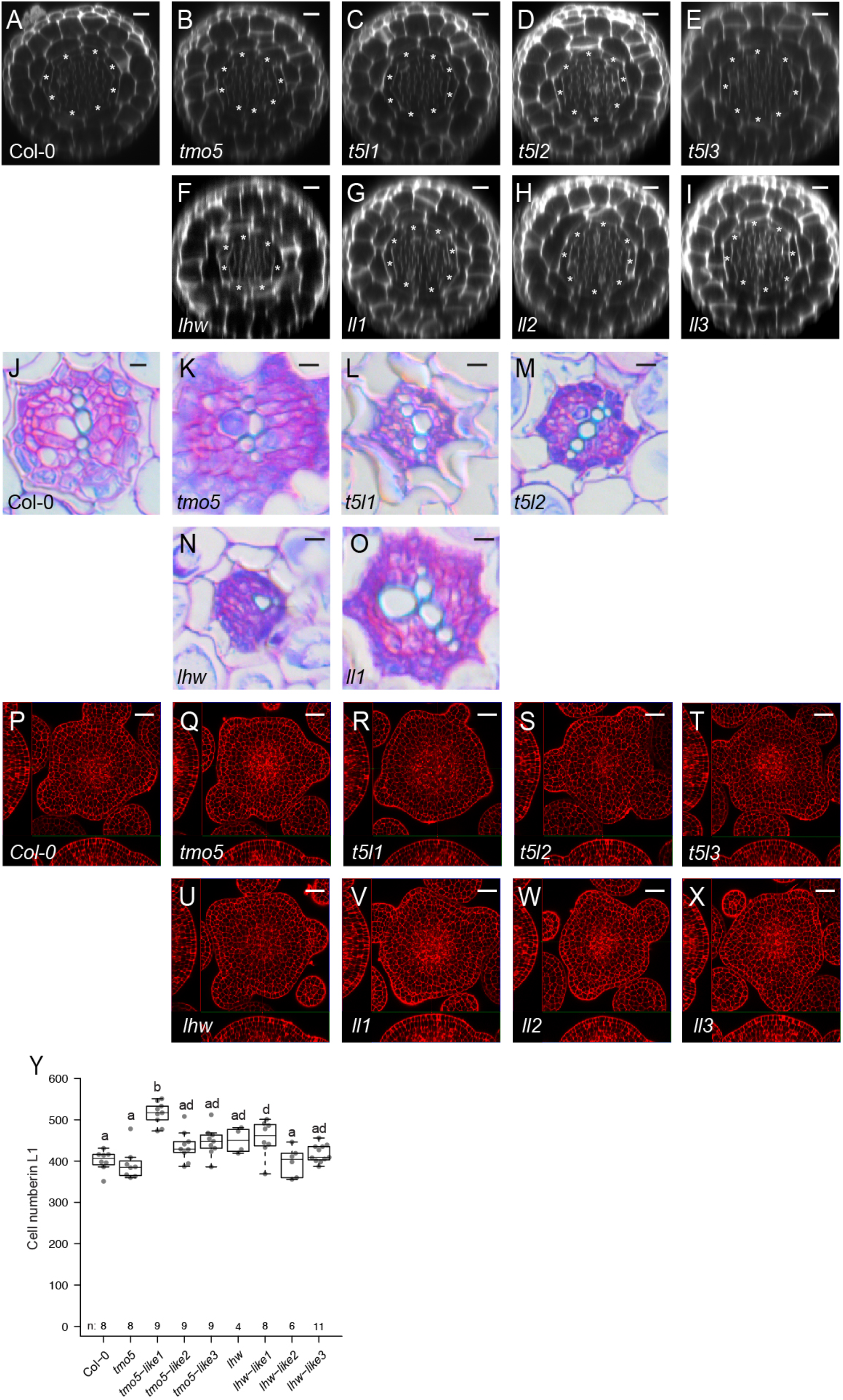
Phenotype of single mutants reveals functional specificity in TMO5 and LHW clade members. Ortho-views of z-stacks confocal images of Col-0*, tmo5*, *t5l1*, *t5l2*, *t5l3*, *lhw*, *ll1*, *ll2*, *ll3* roots. (A-I) Confocal images of 5-day-old root apical meristems stained with MPs-PI, (J-O) 10-day-old root undergoing secondary growth and (P-X) shoot apical meristems. Scale bars: (A-I) 10 µm; (J-Z) 20 µm. (Y) Quantification of cell number in L1 layer of the shoot apical meristems. Lowercase letters indicate significantly different groups as determined using a one-way ANOVA with post-hoc Tukey HSD testing (p ≤ 0.05). Asterisks in A-I and dashed outline in J-O indicate endodermis.

**Supplemental Figure S4.**
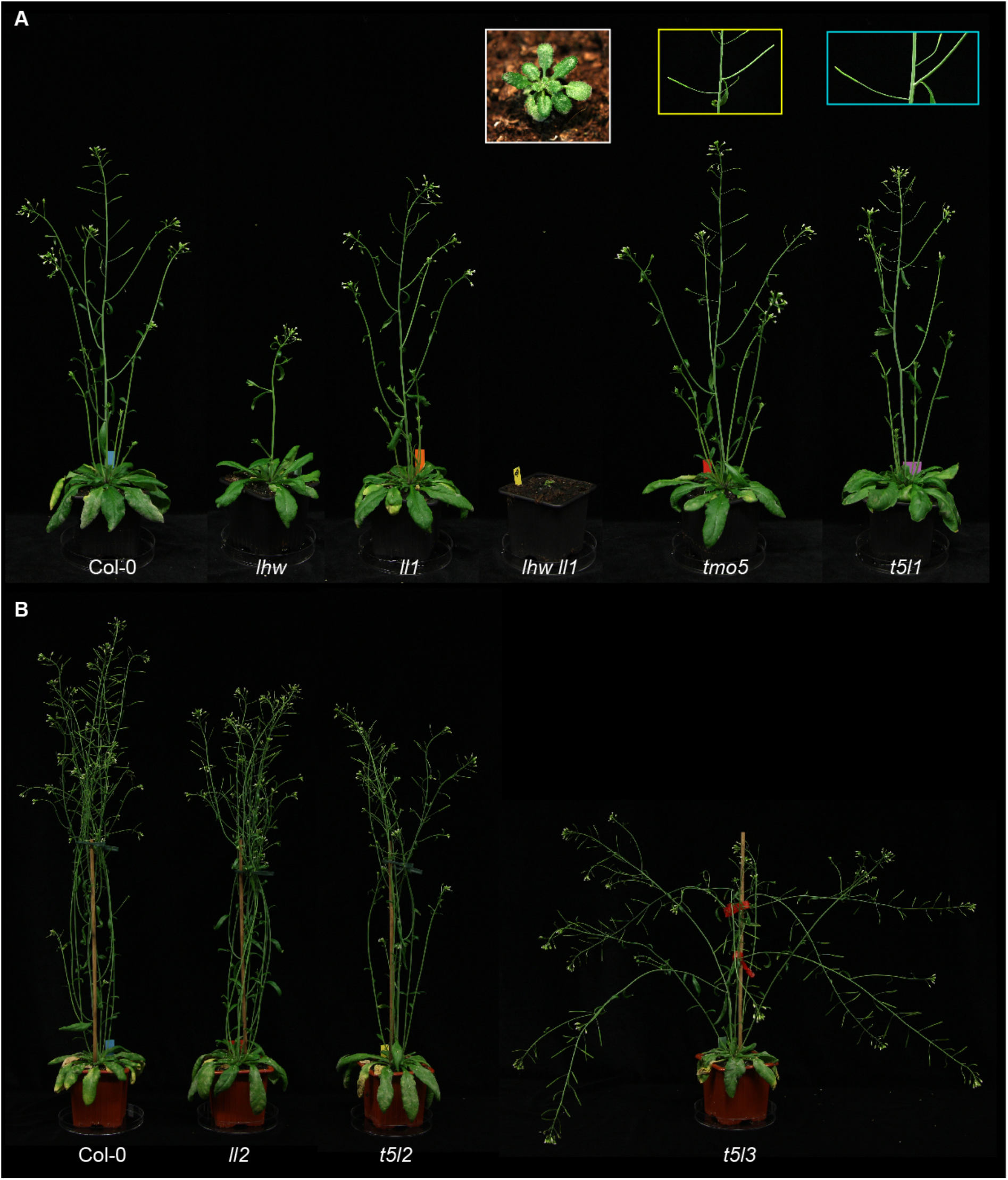
Shoot phenotypes observed in the single mutants of TMO5 and LHW clade members. (A) Shoot phenotype of 38-day-old plants of the indicated genotypes. 50-day-old plant of *lhw ll1* mutant is showed in a white rectangle. Yellow and blue rectangles represent phenotypes of *tmo5* and *t5l1* single mutants, respectively, where silique appears before the last inflorescence branch. (B) Shoot phenotype of 47-day-old plants of the indicated genotypes.

**Supplemental Figure S5.**
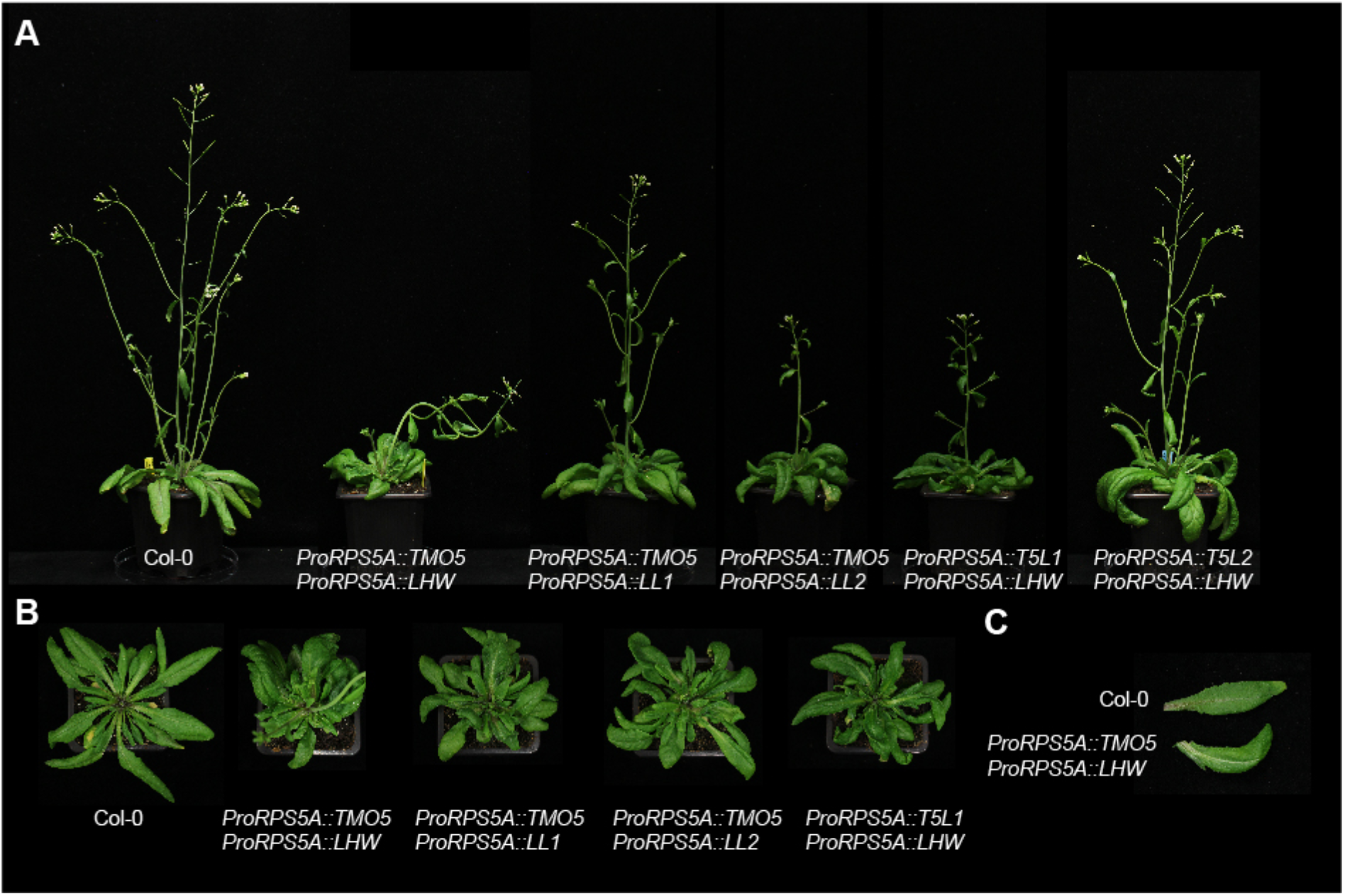
Shoot, rosette, and leaf phenotypes observed in overexpression lines of TMO5 and LHW clade members. (A) Shoot phenotype of 36-day-old plants of the indicated genotypes. (B) Top view of rosettes of the indicated genotypes. (C) Leaf phenotype of *ProRPS5A*::TMO5 x *ProRPS5A*::LHW line.

**Supplemental Figure S6.**
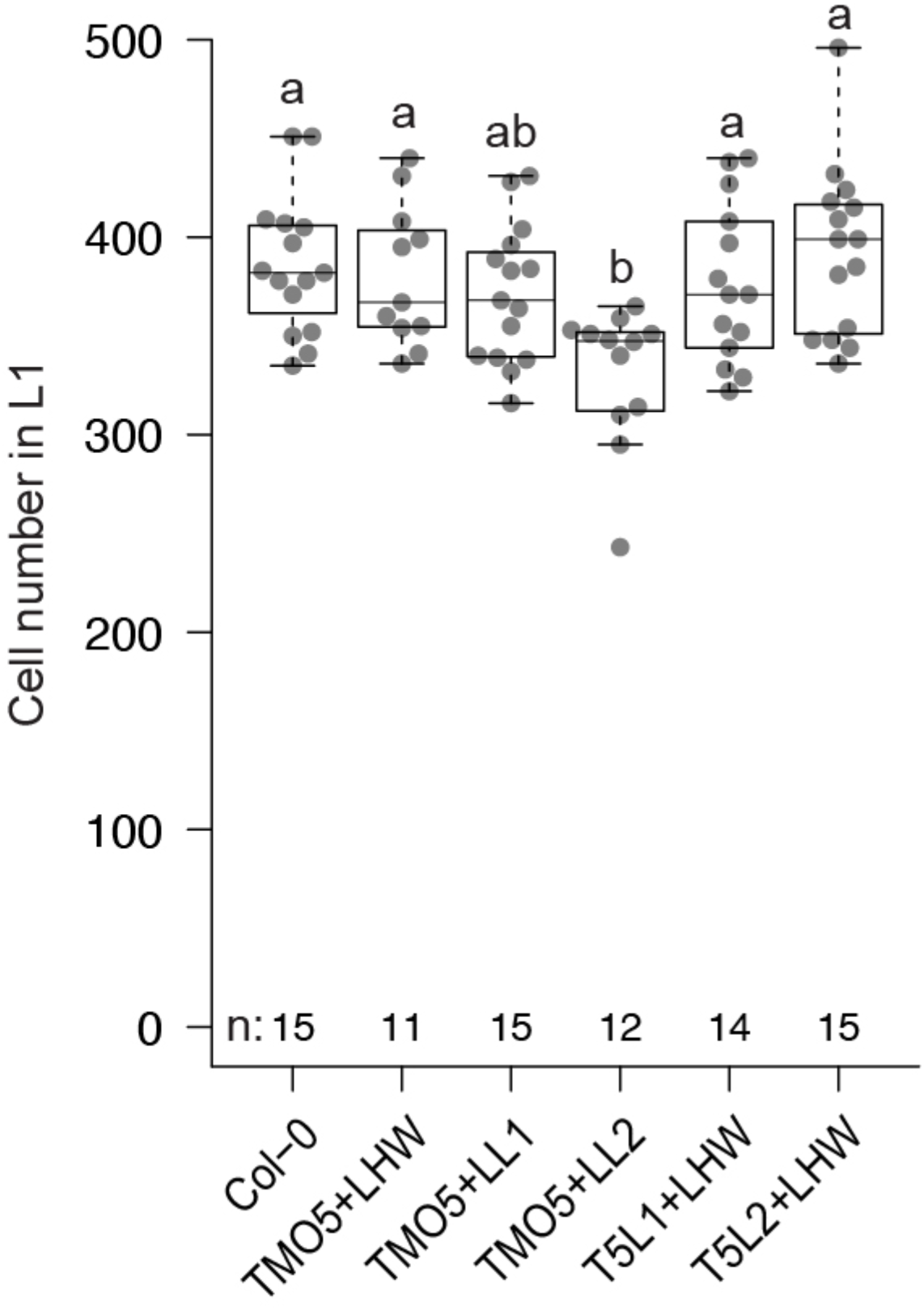
Quantification of cell number in overexpression lines of TMO5 and LHW clade members. Cells were counted in L1 layer of shoot apical meristems. Lowercase letters indicate significantly different groups as determined using a one-way ANOVA with post-hoc Tukey HSD testing (p≤0.05).

**Supplemental Figure S7.**
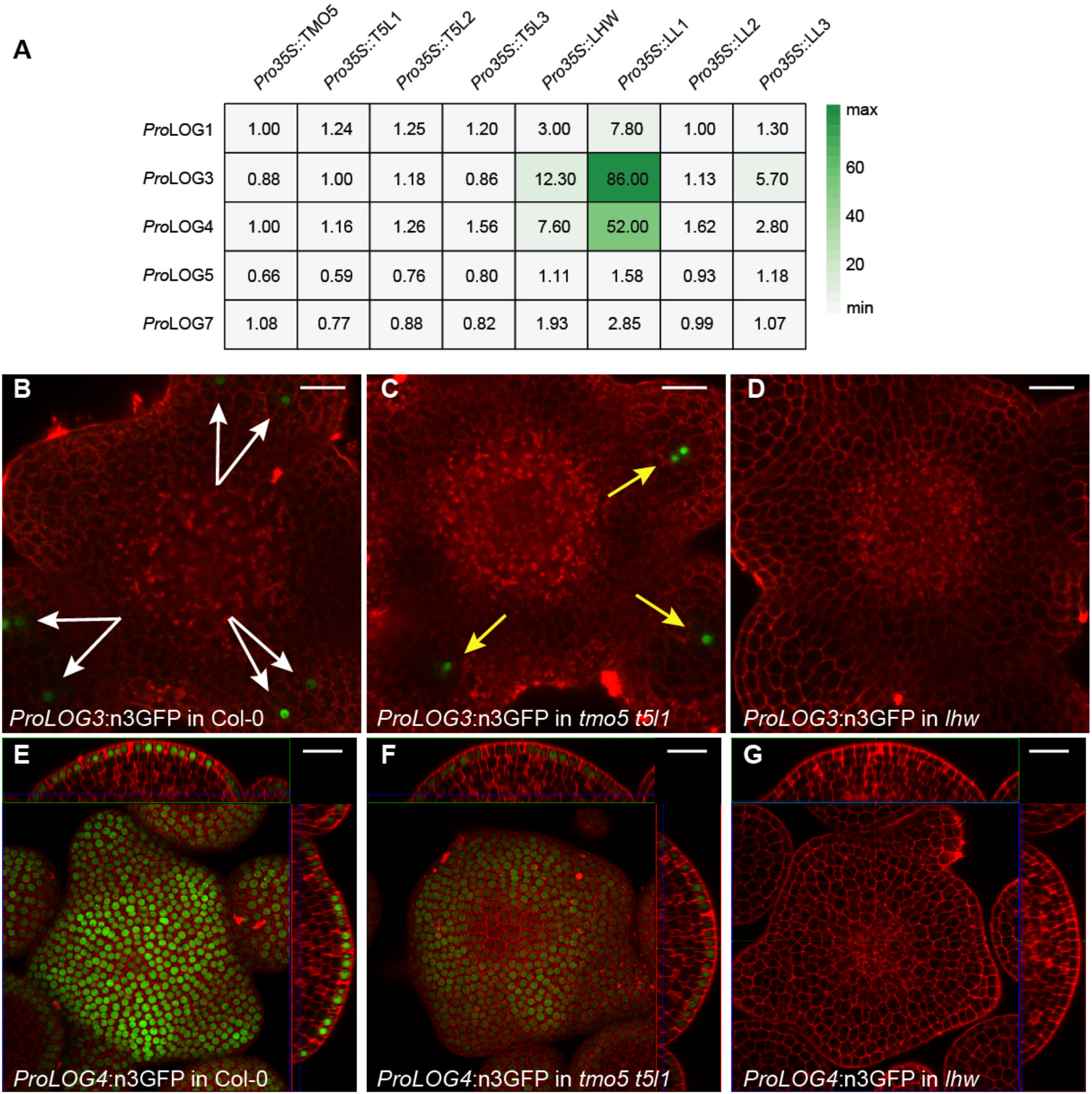
Single TMO5 and LHW clade members affect target gene expression differently. (A) Heat map shows relative changes of gene expression from *LOG* promoters after overexpression of combinations of single TMO5 and LHW clade members in quantitative gene expression assays in *Arabidopsis* protoplasts. Values are FLuc/RLuc ratios, n = 4-6. (B) *ProLOG3* is expressed in two cell files (white arrows) in Col-0 shoot apical meristems compared to (C) a single cell file (yellow arrows) in *tmo5 t5l1* double mutant and (D) undetectable expression in *lhw* mutant. (E) Signal of *ProLOG4* is (F) decreased in *tmo5 t5l1* double mutant and (G) missing in *lhw* mutant. Central squares in E and F represent maximum intensity projection. Scale bars: 20 µm.

**Supplemental Table S1.** Overview of all data and statistics.

**Supplemental Table S2.** List of primers used in this study.

**Supplemental Table S3.** List of plasmids used in this study.

